# Automated cryo-volume EM for high-resolution 3D imaging and *in situ* structural analysis of cells and tissues

**DOI:** 10.64898/2026.06.21.733621

**Authors:** Pavel Křepelka, Jana Moravcová, Zuzana Trebichalská, Elena Buglakova, Lenka Šmerdová, Hana Nedozrálová, Jaroslav Straník, Maria Rosario Fernandez-Fernandez, Pavel Plevka, Anna Kreshuk, Jiří Nováček

## Abstract

Cryo-volume electron microscopy (CVEM) enables three-dimensional imaging of biological ultrastructure in a near-native state but has been limited by low image contrast and charging artifacts that hinder data interpretation and complicate automation of data acquisition. Here we present an experimental and computational workflow that combines orthogonal cryo-SEM imaging, spot-geometry optimized O+ plasma-FIB milling, dedicated acquisition-control routines, and dedicated image alignment procedure. The workflow enables autonomous acquisition of volumetric datasets from vitrified cells and tissues at ∼15–20 nm isotropic resolution. In addition, sub-volume averaging of 113 nuclear pore complexes extracted from CVEM dataset of Cos-7 cell yielded its reconstruction at 9.4 nm resolution. Together, these results establish CVEM as a robust platform for autonomous high-resolution volumetric imaging and structural analysis of vitrified biological specimens.

## Introduction

Cellular function emerges from the spatial organization of biomacromolecules, and understanding the underlying mechanisms is facilitated by three-dimensional imaging at nanometer-scale resolution. Volume electron microscopy (vEM)^1^ has become a central approach for reconstructing cellular and tissue ultrastructure in three dimensions and enables quantitative analysis of complex biological systems across mesoscale dimensions.^2–8^ Among available vEM approaches, focused ion beam–scanning electron microscopy (FIB–SEM)^9,10^ enables automated serial sectioning and imaging of biological specimens with a slice thickness in the 5–50 nm range. The approach has been successfully applied to reconstruction of large biological volumes, including complete neuronal circuits and tissue architectures^3–6^.

Despite the high attainable SEM resolution, the structural information content of conventional vEM remains constrained by specimen preparation procedures involving chemical fixation, dehydration, and heavy-metal staining.^11^ These treatments alter native cellular organization and limit interpretation of molecular-scale architecture. In contrast, cryo-electron tomography (cryo-ET) combined with sub-volume averaging enables *in situ* structural analysis of macromolecular complexes at near-atomic resolution while preserving specimens in a vitrified near-native state.^12–15^ However, cryo-ET is typically restricted to lamellae of ∼100–300 nm thickness, corresponding to only a small fraction of the total cellular volume.

Cryo-volume electron microscopy (CVEM) has recently emerged as a promising strategy to bridge the gap between vEM and cryo-ET by enabling volumetric imaging of unstained vitrified biological specimens.^16–19^ Recent developments, including interleaved scanning strategies^20^ and plasma-FIB milling,^19,21^ substantially improved the feasibility of cryogenic volume imaging. Nevertheless, the number of available high-quality CVEM datasets remains limited. This limitation partly reflects the specific image-formation mechanism in cryo-SEM imaging of vitrified biological samples. Unlike conventional vEM, contrast in CVEM arises predominantly from charge accumulation on non-conductive structures (“charging contrast”).^19,22^ The metastable nature of the created electrostatic potential makes image formation highly sensitive to acquisition conditions and frequently introduces contrast fluctuations, geometric distortions and local charging artifacts. These characteristics complicate autonomous data acquisition and reduce the robustness of standard SEM auto-functions such as focusing, stigmation, and objective lens alignment. In parallel, systematic characterization of CVEM data quality, including attainable resolution, information depth and radiation damage, remains limited, hindering optimization and broader adoption of the method.

Here we present an integrated experimental and computational workflow for autonomous high-resolution CVEM imaging. The workflow combines orthogonal SEM imaging geometry, optimized O+ plasma-FIB milling, dedicated routines for auto-focussing, auto-stigmation, and procedure for tailored image post-processing. We quantitatively characterize the attainable information content of CVEM data and demonstrate near-isotropic structural information at ∼15–20 nm resolution. In addition, we show that CVEM signal support sub-volume averaging of macromolecular assemblies directly within intact vitrified cellular volumes.

## Results

### Orthogonal imaging improves CVEM contrast and resolution

First, we investigated the influence of experimental geometry on CVEM image quality and workflow robustness. The standard through-the-trench geometry (Fig S1), in which the imaging face is tilted by approximately 38° relative to the electron beam, was used as the reference setup. Image quality was evaluated using (I) single-image Fourier ring correlation (siFRC),^23^ (II) image contrast, and (III) charging artifact levels (Fig. 1; see Methods section for details). The measured siFRC_0.143_ resolution ranged between 21 and 28 nm (Fig. 1B, S2B) for the through-the-trench geometry and the average image contrast was ∼18% (Fig. 1A; see Methods for contrast score calculation). We then iteratively varied the imaging angle while maintaining otherwise identical acquisition conditions. Resolution and contrast progressively improved as the imaging face was brought to perpendicular orientation relative to the electron beam (data not shown). Under orthogonal imaging conditions, siFRC_0.143_ resolution improved to 15–19 nm (Fig. 1B, S2B) and image contrast increased to ∼42% (Fig. 1A), corresponding to more than a twofold improvement relative to the through-the-trench geometry.

**Figure 1:**
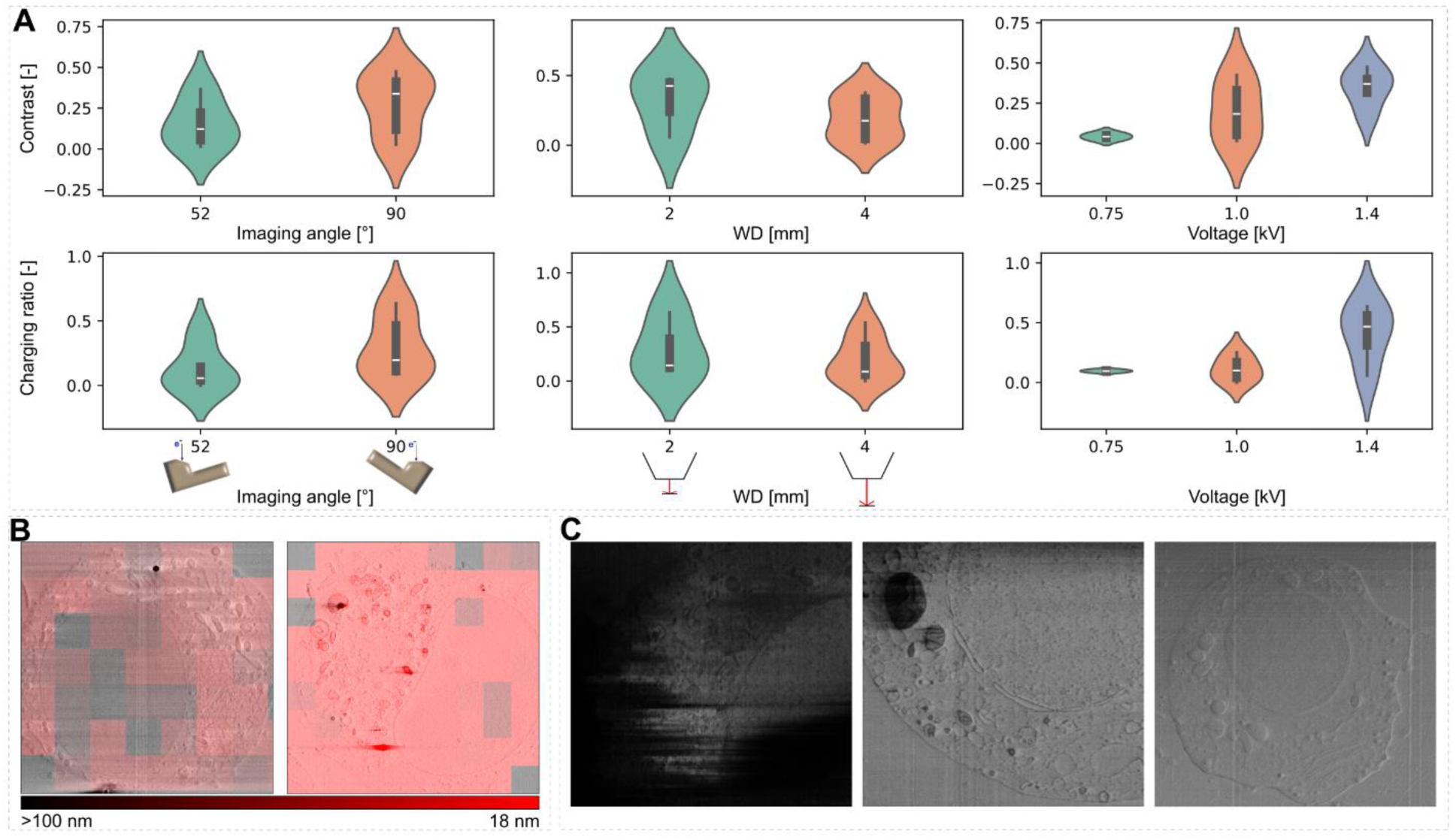
Optimization of cryo-SEM image quality. The overall image contrast and level of charging artifact observed in the cryo-SEM images measured for the through-the-trench and orthogonal geometry (A, *left,* 1 kV, 25pA). While the level of charging artifacts is similar for the two geometries, the observed image contrast increases significantly for the orthogonal geometry. The sample distance from the objective lens pole piece does not influence the level of charging artifacts, but shorter distances significantly improve the image contrast (A, *center*). Image contrast increases with the electron energy which level of charging artifacts remains constant below 1.0 kV, but starts to significantly increase for higher acceleration voltages (A, *right*). A comparison of siFRC_0.143_ values calculated across 256×256 pixel patches in cryo-SEM images for optimal conditions in case of through-the-trench (1.4kV, B, *left*) and orthogonal geometry (1.0kV, B, *right*). A comparison of cryo-SEM images demonstrating high-contrast and low level of charging artifacts (C, *center*, 1 kV, 25 pA, 90°); high level of charging artifacts observed at increased acceleration voltage (C, left, 1.4kV, 25pA, 90°), and decrease in image contrast observed for through-the-trench geometry (C, right, 1.0kV, 25pA, 52°).

We further observed that primary electron beam energy strongly affects both image contrast and charging artifact intensity. Increasing the beam energy enhanced contrast but also increased charging artifacts (Fig. 1A). Electron energies above 1 keV resulted in substantial charging under orthogonal imaging conditions, whereas ∼1 keV provided the optimal balance between contrast and charging stability (Fig. 1A). In the conventional through-the-trench geometry, reaching high contrast required working at higher accelerating voltages (1.2–1.5 keV), which further increased susceptibility to overcharging. In addition to imaging angle, image quality improved when reducing the working distance from the eucentric position (4 mm) to 2 mm from the pole piece (Fig. 1A). This effect was observed consistently across all tested electron beam energies.

### Optimized CVEM geometry enables stable orthogonal imaging

The improved performance of orthogonal imaging motivated the development of an updated CVEM data acquisition workflow optimized for stable perpendicular imaging and milling (Fig. 2A). In the first step, the specimen surface is polished at a shallow milling angle of 3° to remove contamination and surface irregularities. The sample is subsequently rotated by 90°, and an initial trench is milled perpendicular to the polishing direction. This setup minimizes residual curtaining artifacts because the final imaging face is created perpendicular to the direction of the milling artifacts on the face prepare in previous step. A protective organometallic platinum layer is then deposited onto the cleaned surface, followed by preparation of the imaging face at a 16° milling angle. During acquisition, the microscope stage alternates between perpendicular imaging and perpendicular milling geometry, maintaining 90° beam-to-sample orientation for both electron imaging and ion milling. Sample tilting and subsequent stabilization prolonged the acquisition of a single slice. Combined with need for more frequent call of auto-functions for image centering and focusing, single milling-imaging cycle of the CVEM workflow presented here is ∼18 seconds longer (∼7.5%) compared to the through-the-trench approach while enabling stable milling and high-quality imaging (Fig. 2B, S2A).

**Figure 2:**
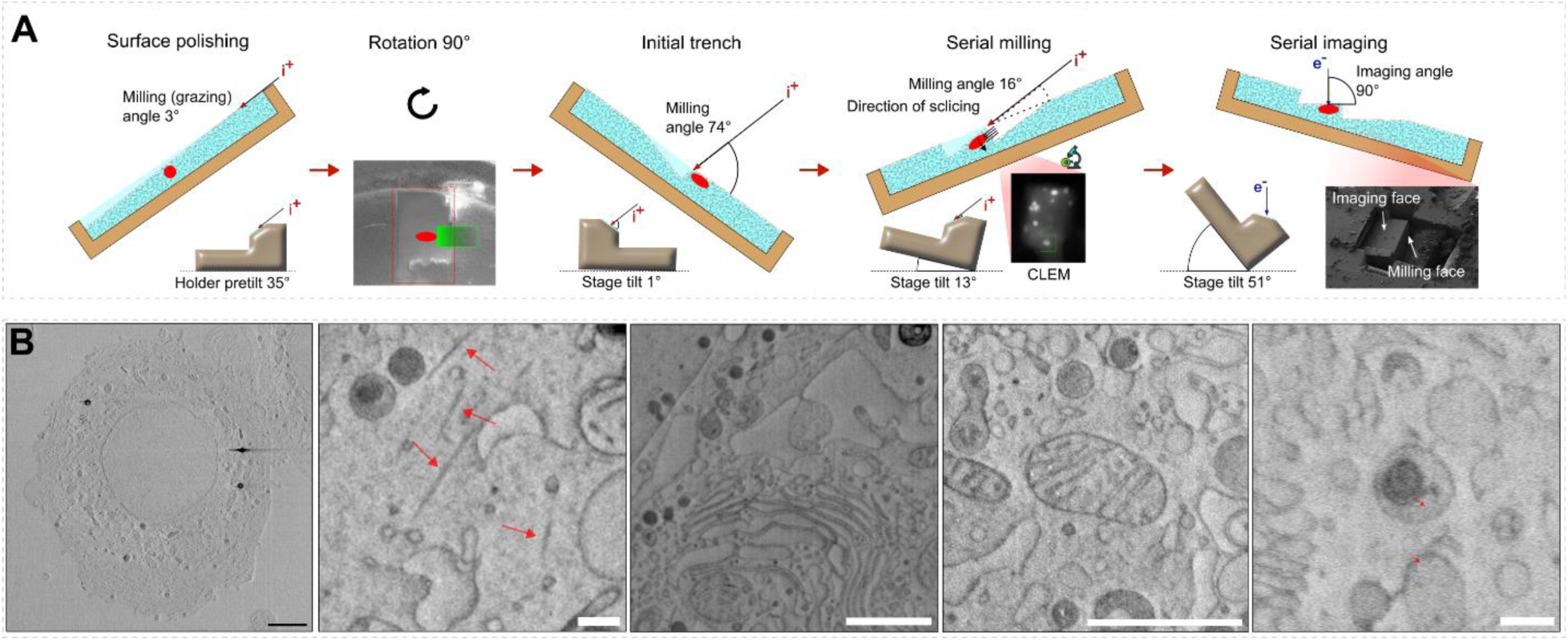
A graphical representation of the FIB/SEM data acquisition workflow developed in this work (A). First, the sample surface is polished by high current and low milling angle. Then the sample is rotated by 90°. The rotation ensures the milling of the initial trench perpendicularly to “curtains” created during previous step. Sample is ablated through the polished plane of the initial trench sputter coated with organometallic platinum layer from GIS. The sample is tilted between milling and imaging to allow for perpendicular imaging. Overview image of INS-1E cell and details highlighting tentative actin filaments (red arrows), Golgi apparatus, mitochondrion, and insulin granule (B). The scale bars correspond to 1 μm (black) and 200 nm (white), respectively.

### Low-energy imaging preserves near-isotropic structural information

We investigated how electron beam current and imaging dose influence image quality and sample integrity. Images were acquired at constant total electron dose (0.2 e−/Å²) using 1 keV electrons under orthogonal imaging conditions. Contrast remained stable across beam currents between 1 and 100 pA, whereas siFRC resolution gradually deteriorated at higher beam currents (Fig. 3A). Considering both image quality and acquisition speed, beam currents between 6 and 25 pA were selected as optimal operating conditions.

**Figure 3:**
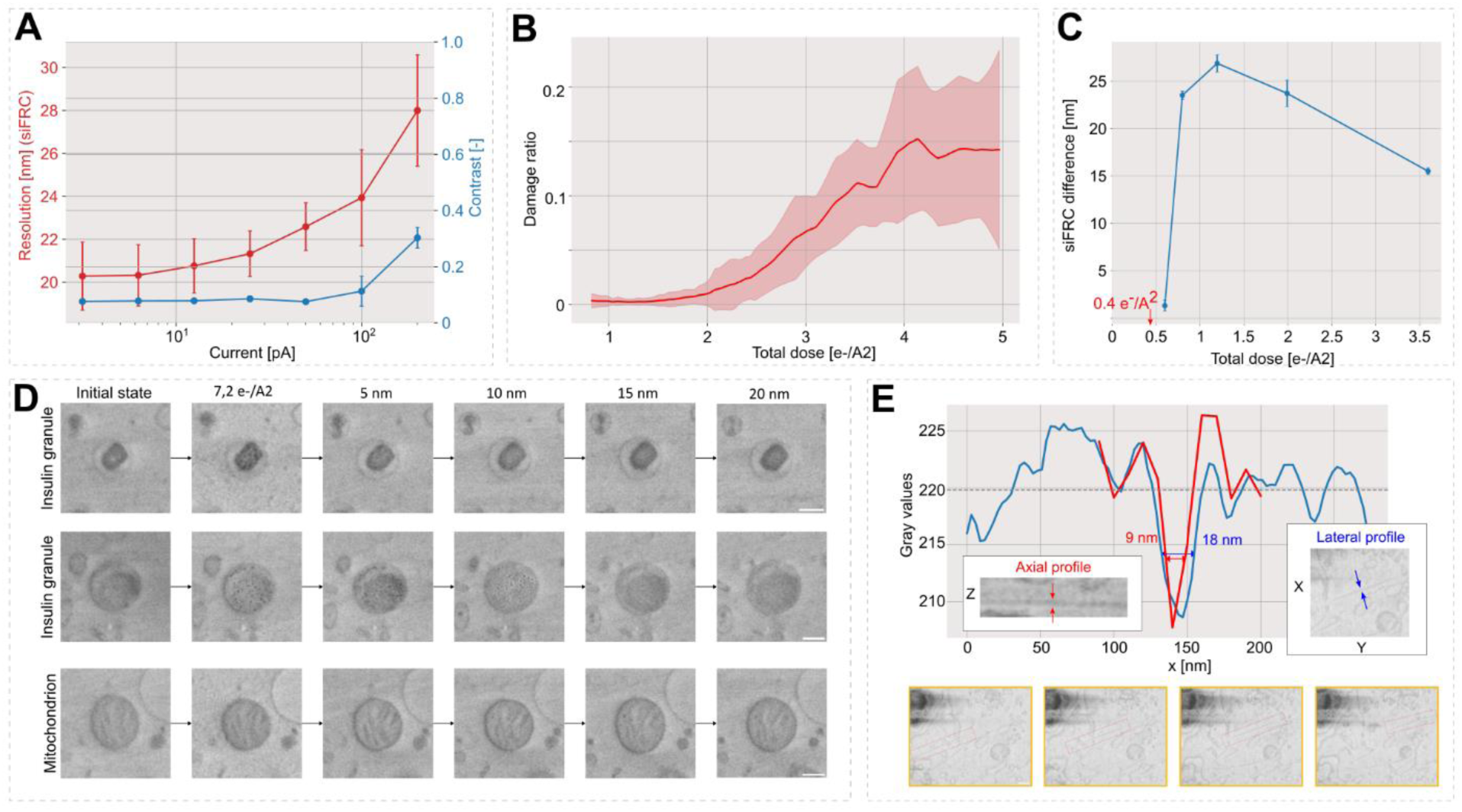
Combined graph showing the siFRC_0.5_ and contrast dependence on the imaging current (A). The total dose deposited to the sample was constant for images acquired under with electron beam currents. The contrast in the images acquire at 1 kV is constant for electron beam currents small than 100 pA. The attainable resolution show approximately log-squared dependence on the imaging current. Level of radiation-induced structural changes of the sample (“dark” spots in the data) observed under eight different imaging conditions. The data was acquired under following conditions: (1) U=1 kV, I=13 pA, LI=255, DT=25 ns, PS=5 nm, HFW=30 μm; (2) U=1 kV, I=13 pA, LI=255, DT=25 ns, PS=2.5 nm, HFW=30 μm; (3) U=1 kV, I=13 pA, LI=32, DT=25 ns, PS=1.25 nm, HFW=30 μm; (4) U=1 kV, I=13 pA, LI=255, DT=100 ns, PS=5 nm, HFW=30 μm; (5) U=1 kV, I=13 pA, LI=64, DT=25 ns, PS=5 nm, HFW=10 μm; (6) U=1 kV, I=13 pA, LI=255, DT=25 ns, PS=5 nm, HFW=10 μm; (7) U=1 kV, I=13 pA, LI=255, DT=25 ns, PS=5 nm, HFW=10 μm; (8) U=2 kV, I=13 pA, I=255, DT=25 ns, PS=5 nm, HFW=10 μm, where U is the accelerating voltage, I is the imaging current, LI is the number of lines integrated during scanning, DT is dwell time, PS is pixel spacing, and HFW corresponds to the width of scanned area. SiFRC resolution change measured in 5 nm sample depth as a function of electron dose applied to the sample (C). The resolution difference is calculated as a difference of the mean siFRC_0.143_ measured on the face before and after imaging with a defined dose. The 5 nm of the sample was ablated before the acquisition of the second image and both micrographs were acquired with 0.4e/Å^2^. Differential sensitivity of cellular ultrastructures to the 1 kV electron beam imaging and sample depth (D). A cell was irradiated with 7.2 e/Å² followed by sequential ablation of 5 nm section and imaging with 0.4 e/Å². Insulin granules (D, two lines at the top) show high sensitivity to the electron beam irradiation and structural changes can be observed up to 15 nm below the surface subjected to 7.2 e/Å² dose. No structural changes were observed for mitochondrion exposed to the same irradiation (D, bottom line) already at 5 nm below sample surface. Lateral and axial profiles of a filamentous structure identified in CVEM data and a series of raw images collected with 10nm slice distance (E). A sequence of four subsequent micrographs from CVEM dataset focused on the region with the filamentous structure is shown at the bottom.

To evaluate electron-induced sample damage during low-energy SEM imaging, we used INS-1E cells containing crystalline insulin granules^24^ as an *in cellulo* reporter highly sensitive to electron beam damage (Fig 3B--D). Structural alterations were quantified from dose-series experiments by segmenting dark contrast features associated with sample alteration (Fig 3B). For insulin granules, structural changes became apparent at cumulative doses exceeding ∼0.6 e−/Å², while lower doses preserved ultrastructural integrity. Consequently, the imaging dose used for volumetric acquisition was conservatively limited to 0.4 e−/Å² (Fig 3C). Notably, individual ultrastructures exhibited varying degree of sensitivity to the electron beam. For instance, mitochondria have not demonstrated any discernible global structural alterations even after a cumulative dose of 7.2 e/Å^2^ (11.6e^-3^ C/m^2^; Fig 3D).

Further, we analysed the electron-sample interaction depth during electron imaging. Monte Carlo simulations of low-energy electron interactions with vitreous ice demonstrated that the majority of primary interactions occur within the first 10 nm below the sample surface (Fig S3). At 1 keV, only 7.6% of primary electrons were backscattered and contributed to secondary - electron generation, while 92.4% of SE-I generating interactions occurred within the first 10 nm of the specimen. In contrast, simulations at 5 keV predicted substantially deeper penetration and broader signal delocalization (Fig. S3). Experimental analysis of dose-dependent structural deterioration confirmed that no detectable ultrastructural damage occurred below ∼15 nm sample depth. For that, the sample face has been imaged with 7.2 e/Å^2^ and 5 nm sections were gradually ablated, and the effect on different ultrastructures was evaluated from images collected with 0.4 e/Å^2^ (Figs. 3D, S4). No observable artefacts from structural changes were observed above 15 nm sample depth. Subsequently, the resolution deterioration in a 5 nm sample depth was then measured after deposition of 0.4 – 7.2 e/Å^2^ (Fig. 3C). The results show that cumulative dose above 0.6 e/Å^2^ resulted in a systematic decrease of the maximal resolution. Finally, we analysed filamentous structures acquired with 10 nm slice spacing (Fig. 3E) to experimentally validate axial information depth. The measured full width at half maximum (FWHM) values were 18 nm laterally and 9 nm axially, indicating that detectable secondary - electron signal originates from less than ∼10 nm below the surface.

### Plasma ion milling improves imaging quality and workflow efficiency

We compared different FIB ion sources with respect to (I) milling efficiency, (II) image quality, and (III) surface artifact formation (Fig. 4C). Milling rates were measured separately for vitrified ice and an organometallic platinum protective layer using Ga, Xe, Ar, O and N ions. For organometallic platinum, milling rates correlated with the atomic number of the ion species (Fig S5). In vitrified ice, however, oxygen and nitrogen plasma exhibited substantially higher milling efficiency than expected based solely on ion mass. The Xe+/O+ milling rate ratio decreased from 3.3 for platinum precursor to 1.7 for vitrified ice while the Ga+/O+ ratio changed from 2.3 to 0.86. These observations suggest that oxygen plasma milling benefits from a combination of physical sputtering and chemical etching mechanisms. On average, we have observed ∼20x higher milling rates for vitrified ice compared to platinum milling (Fig. S5).

**Figure 4:**
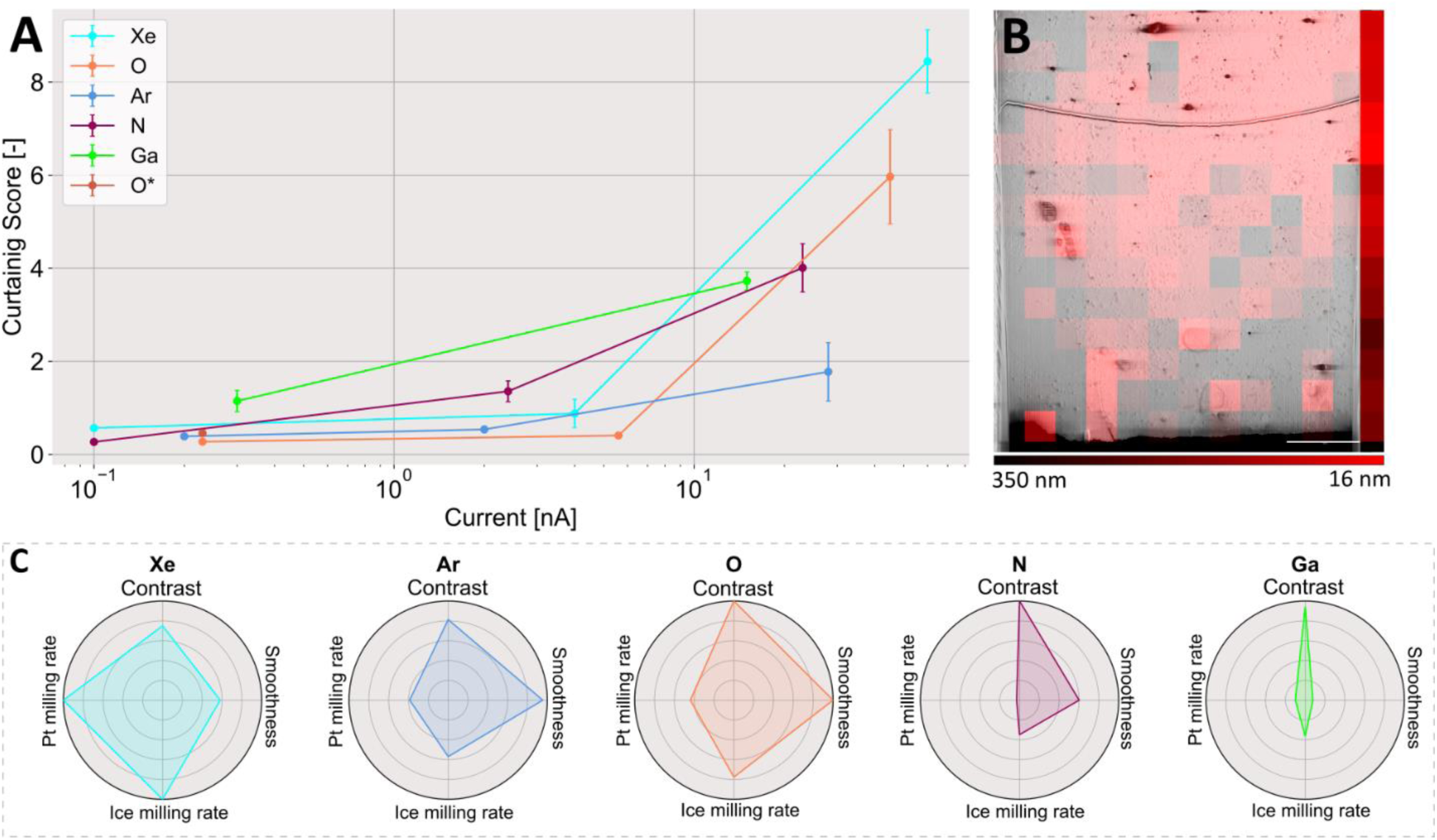
Curtaining score (higher score corresponds higher level of curtaining artifacts) dependence on the milling current for respective ion beam sources (A). Local resolution distribution across the depth of the imaging face for orthogonal milling with O^+^ (610 pA; B). The regions of the image with siFRC_0.143_ resolution better than 18 nm is highlighted in red. The stripe on the left side shows average resolution at different depth of the imaging face. Data quality (siFRC_0.143_ < 18 nm) is preserved to approximately 20 μm distance from image face front. The scale bar corresponds to 1 μm. Radar plots comparing cryo-SEM image contrast, milling rate into organometallic precursor and vitrified ice, and smoothness of the imaging face for individual focused ion beam sources (C). Larger area highlighted in the radar plot corresponds to better performance of the ion source.

Curtaining artifacts were quantified using Fourier-based analysis of surface irregularities^19^ (Fig 4A, S5B; see Methods). Consistent with previous reports,^19^ heavier ions generally increased curtaining intensity. Oxygen plasma represented an important exception because multiple oxygen ion species induced beam asymmetry and spot distortions that increased surface artifacts. To address this limitation, we developed a dedicated alignment procedure for spot-shape optimization using the auxiliary immersion lens of the SEM column to correct beam geometry (Methods, Fig. S5D). Following the application of this alignment procedure, O+ plasma milling produced the lowest curtaining levels among all tested ion sources for beam currents below 10 nA (Fig 4A).

Argon plasma exhibited comparable milling quality and represents a suitable alternative for systems lacking immersion-mode beam compensation (Fig 4A). Xenon plasma maintained acceptable surface quality at lower currents but exhibited substantially increased curtaining at higher currents. To complement the data for different plasma sources, surface roughness was also evaluated across the depth of the polished face (Fig. 4B, data shown for milling with 610 pA O^+^ plasma). The ultimate data quality (siFRC_0.143_<18 nm) was systematically observed to ∼20 um face depth (for the range of currents defined below), above which the average resolution gradually deteriorated. Overall, plasma currents between 0.5 and 1 nA provided the optimal balance between milling speed (Fig. S5A,C) and surface quality for CVEM imaging (Fig. 4B,C).

### Autonomous acquisition and deformation correction improve workflow robustness and data interpretation

Low signal-to-noise ratios and charging-induced instabilities place specific requirements on CVEM automation. Standard SEM autofocus procedures which search for maximal image contrast frequently failed under cryogenic conditions and acquisition of image series often introduced unnecessary radiation exposure. To address this limitation, we implemented and further developed line-based optimization algorithms that determine focusing or stigmation or objective lens alignment parameters within a single scan^25^ (Fig 5A). During acquisition, the optimized variable (e.g. working distance) is continuously varied across scanned lines, while image-quality metrics based on power spectral density in middle frequencies are evaluated in real time (Fig 5A). The optimal parameter value is subsequently extracted from the resulting focus profile. The workflow further incorporates continuous siFRC-based image quality monitoring and automatic activation of optimization routines when resolution falls below a predefined threshold (Fig 5B). These procedures were integrated together with workflow management, drift correction and acquisition control into the open-source software FIB-SEM Maestro (https://github.com/cemcof/FIBSEM_Maestro).

**Figure 5:**
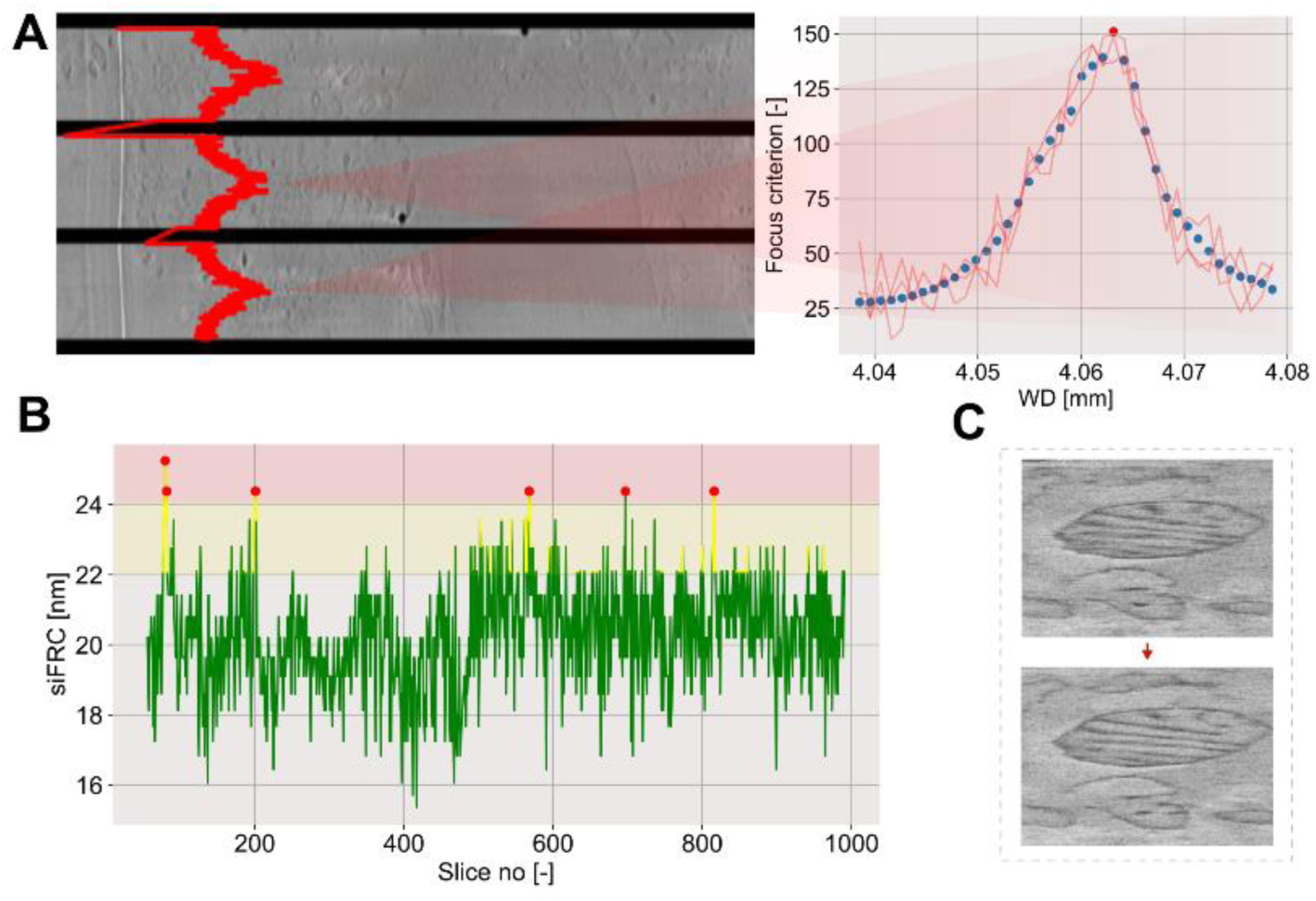
The implemented routine for autofocussing calculates focus criterion from a single line scan while changing the focus distance (working distance) between subsequently scanned lines (A). The siFRC_0.143_ resolution is measured for the selected region of the image in real-time and used to trigger initiation of the auto-fuctions once the resolution drops below predefined threshold (B). The x-y cross-section of mitochondrion from CVEM dataset acquired on cos-7 cell (C). The original image displays line-to-line misalignment (C, top). The same mitochondria after application of the deformation correction algorithm presented in this study (C, bottom).

Dynamic charging nature of CVEM imaging introduces additional deformation, which, in combination with low signal-to-noise, is challenging to mitigate using standard vEM data alignment algorithms.^26^ CVEM data exhibit correlated line-to-line distortions caused by local charging fluctuations (Fig. 5C). We developed a dedicated deformable alignment algorithm that models image distortions as smooth displacement fields varying between scanned lines. Pairwise cross-correlation between consecutive slices was used to recover true line positions, and the resulting displacement fields were applied to raw images. The correction substantially improved interpretation of cellular ultrastructure and facilitated subsequent segmentation and structural analysis (Fig 5C).

### High-resolution CVEM enables analysis of cellular architecture and macromolecular assemblies

The developed workflow was used for autonomous acquisition of volumetric datasets across six vitrified specimen types, including mammalian cells, muscle tissue, and bacteriophage-infected *Staphylococcus aureus* biofilms (Fig 6). CVEM datasets of whole mammalian cells resolved ultrastructures observable in vEM such as mitochondria (including clearly defined membrane separation in cristae), Golgi apparatus, endoplasmic reticulum, nuclear membrane, but more importantly, also structures which are more challenging to observe in vEM data such insulin grantules and proteinaceous filamentous structures (Fig. 2B,6). Therefore, CVEM data enable visualization and measurement of membrane-contact sites between mitochondria, endoplasmic reticulum and the nucleus under near-native conditions (Fig.6C). Notably, we identified helical tau filaments in HEK293T Tau RD cells and directly measured their width and helical pitch within the cellular environment (Fig. 6D). In muscle tissue, the workflow enabled reconstruction of mitochondrial networks and sarcomeric Z-discs across large cellular volumes (Fig. 6F).To evaluate whether coherent structural information can be extracted from CVEM datasets, we performed sub-volume averaging of nuclear pore complexes (NPCs) identified in COS-7 cells (Fig. 6E). A total of 239 NPC particles were initially selected manually and subjected to iterative alignment and 3D classification using in-house developed pipeline implemented in Python. Following multiple rounds of 3D classification, 113 NPCs were selected for final refinement using gold-standard Fourier shell correlation protocol, which resulted in 9.4 nm resolution (FSC_0.143_) CVEM model.

**Figure 6:**
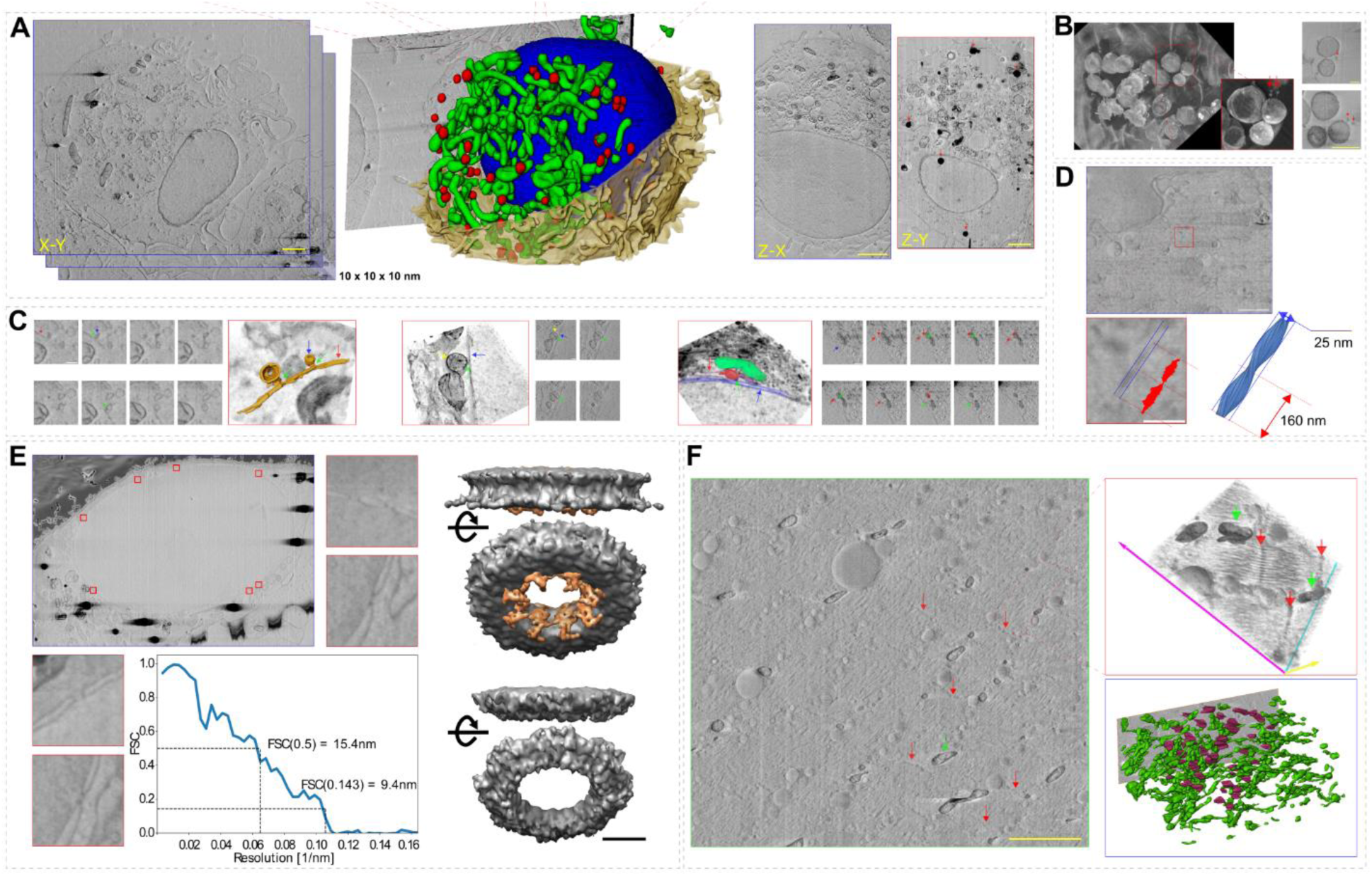
Triplanar views of the entire Cos-7 cell dataset together with the 3D segmentation of mitochondria (green), lipid droplets (red), and nucleus (blue). Red arrows in Z–Y projections indicate charging artifacts from lipid droplets (A). The φ812 bacteriophage (red arrows) infected *S. aureus* biofilm CVEM data showing bacteriophages infecting the bacteria (3D maximum-intensity projection – left and individual slices – right; B). Membrane contacts with cytoskeleton or organelles: actin (red)–ER (blue, left), mitochondrion (yellow)–nuclear membrane (blue, middle), mitochondrion (green) – ER (red) – nucleus (blue) contacts (red, right; C). Cross-section of a HEK293 cell showing helical tau filaments. The helical pitch and filament width were measured directly in the cellular context (D). Nuclear pore complexes in Cos-7 cells and sub-volume averaging of CVEM data. Individual pores are highlighted in red frames. The CVEM density of nuclear pore refined using 113 NPC volumes with the corresponding FSC curve from (E). Muscle tissue slice showing Z-discs (red) and mitochondria (green). The top-right inset contains a 3D visualization of the sarcomere. CVEM data of mitochondrial network (green) and Z-discs (maroon) in the muscle tissue are shown in the bottom panel (F). Scalebars: 200 nm (white), 1000 nm (yellow).

## Discussion

The workflow presented here (Fig. 2A) establishes CVEM as a robust and autonomous platform for high-resolution volumetric imaging of vitrified biological specimens. A central feature of the approach is the implementation of orthogonal SEM imaging geometry, which substantially improves image contrast (Fig. 1A), attainable resolution (Fig. 1B, S2B) and workflow stability (Fig. 5) compared to conventional through-the-trench acquisition. Yet, low contrast remains an inherent limitation of all cryo-EM methods. In cryo-TEM, the development of dedicated direct electron detectors^27^ transformed attainable resolution and enabled the “resolution revolution”.^28^ In contrast, CVEM still relies predominantly on general-purpose SEM detectors that were not designed specifically for low-dose cryogenic biological imaging. The orthogonal imaging strategy implemented here substantially improves signal sensitivity and image contrast by reducing signal delocalization and enhancing secondary-electron collection efficiency, which is caused by maximizing the difference between the SE escape depth and the interaction volume.^29,30^ In addition, orthogonal imaging reduces topographic contributions to the detected signal and improves compatibility with immersion-mode SEM imaging. The workflow presented here mitigates these effects by combining shallow-angle milling with perpendicular imaging and milling geometry. Although stage tilting increases acquisition time of single milling-imaging cycle (<10%), the resulting improvements in image quality and robustness substantially outweigh the throughput penalty. The experimental geometry presented here also naturally supports future integration with correlative cryogenic light-electron microscopy workflows (cryo-CLEM). The shallow-angle preparation strategy provides direct optical access to the imaging face and is compatible with integrated fluorescence microscopes available on modern cryo-FIB/SEM platforms. Combined with recent developments in AI-assisted segmentation,^31–34^ automated targeting^35^ and multimodal imaging,^36^ these advances create opportunities for large-scale volumetric structural analysis directly within intact vitrified specimens.

The image quality obtained under orthogonal imaging conditions enables operation at lower accelerating voltages (∼1 keV), which partially suppresses charging artifacts while preserving high contrast. Notably, the workflow is fully compatible with recently developed interleaved scanning approaches,^20^ and its application will further mitigate presence of charging artifacts. Importantly, low-energy imaging also limits the penetration depth of primary electrons and thereby reduces signal delocalization along the z-direction. Through a combination of Monte Carlo simulations (Fig. S3), cumulative dose experiments (Fig. 3B,C), and direct analysis of filamentous structures (Fig. 3E), we demonstrate that detectable structural information originates from less than ∼10 nm below the sample surface under. Together with the experimentally measured lateral resolution (Fig. 1B, S2B), these findings indicate that CVEM can provide near-isotropic structural information at ∼15–20 nm resolution across large cellular volumes. This represents an important distinction from conventional volume EM, where anisotropic information content is frequently introduced by both section thickness and signal delocalization.^37,38^

Our data further highlights the importance of optimized plasma-FIB milling for high-resolution cryogenic volume imaging (Fig. 4C). While noble-gas plasma sources such as argon and xenon are widely used because of their milling stability and reduced curtaining,^21^ we show that oxygen plasma offers additional advantages for vitrified biological materials (Fig. 2B). Oxygen plasma produced unexpectedly high milling rates in vitrified ice compared to organometallic platinum, exceeding expectations based solely on ion mass (Fig S5). These observations are consistent with previous reports describing enhanced oxygen-mediated sputtering of carbon-rich materials^39^ and suggest that a chemical component contributes to ablation in addition to collision-driven sputtering. Reactive oxygen species likely reduce redeposition of sputtered material and thereby improve surface quality and image contrast.^40^ Importantly, however, oxygen plasma produced superior results only after correcting beam-spot asymmetry using the dedicated alignment procedure introduced here (Fig. S5). Following spot-shape compensation, O+ plasma generated the lowest curtaining levels among all tested ion species for currents below ∼10 nA.

A major bottleneck in CVEM has been the instability of conventional SEM automation routines under cryogenic imaging conditions. Standard autofocus and auto-stigmation procedures typically rely on repeated acquisition of multiple images while maximizing contrast-based metrics. Under low-dose cryogenic conditions, these approaches frequently failed and often resulted in unnecessary radiation exposure. The line-based optimization procedures implemented here substantially improve the robustness of autonomous acquisition by determining optimal imaging parameters within a single scan. In parallel, continuous siFRC-based monitoring enables real-time quality control during long-term volumetric acquisition (Fig. 5). These routines were integrated together with drift correction and workflow management into the open-source software FIB-SEM Maestro, enabling stable autonomous acquisition of large volumetric datasets. In addition to low signal-to-noise ratio, CVEM datasets exhibit charging-induced geometric distortions (Fig. 5C) that are difficult to correct using standard volume-EM alignment approaches. We observed that line displacements within individual SEM scans remain correlated within a single image while varying independently between subsequent z-frames. This reproducible behavior enabled development of a dedicated deformable alignment strategy based on line-wise displacement correction (Fig. 5C). The resulting correction improves interpretability of cellular ultrastructure and facilitates downstream segmentation and quantitative analysis.

The CVEM workflow developed here provides structural information extending beyond membrane ultrastructure, which has typically dominated contrast in previously reported CVEM datasets. While membranes and membrane interfaces remain readily identifiable at a level comparable to conventional vEM, the improved information content also enables direct visualization and tracing of proteinaceous structures, including filamentous assemblies and cytoskeletal interactions, within intact vitrified cellular environments. Furthermore, the sub-10 nm reconstruction of the nuclear pore complex demonstrates that coherent structural information can be further enhanced through sub-volume averaging of CVEM datasets acquired using secondary electron imaging. This observation is consistent with recent developments in materials science, where secondary-electron imaging has enabled structural analysis at substantially higher resolution,^41^ highlighting the potential of further advances in CVEM methodology and data analysis.

In conclusion, we present a robust and autonomous CVEM workflow capable of imaging vitrified biological specimens at ∼15–20 nm isotropic resolution. The workflow combines orthogonal cryo-SEM imaging, optimized plasma-FIB milling, quantitative characterization of imaging conditions, and dedicated acquisition-control and deformation-correction algorithms. Together, these developments substantially improve CVEM data quality, acquisition robustness and attainable structural information. Importantly, the presented approach extends CVEM beyond volumetric ultrastructural imaging toward high-resolution structural analysis within intact cellular volumes. Although future developments in imaging strategies, secondary-electron detection and acquisition throughput will likely further improve the attainable information content of CVEM datasets, our findings suggest that CVEM has the potential to complement cryo-electron tomography by enabling structural analysis of macromolecular assemblies across substantially larger cellular volumes while preserving specimens in a near-native state.

## Materials and Methods

### INS-1E culture cultivation

INS-1E cells (AddexBio, cat. no. C0018009) were cultured to 80–90% confluency at 37°C and 5% CO₂ in Advanced RPMI medium (Thermo Scientific™, cat. no. 12633012) supplemented with 10% FBS (heat-inactivated for 30 min at 56°C), 10 mM HEPES (from a 1 M stock solution), 1 mM L-glutamine (from a 200 mM stock solution), 1% penicillin/streptomycin (5,000 units/mL penicillin and 5,000 μg/mL streptomycin), and 3.5×10⁻⁴% β-mercaptoethanol. Cells were dissociated into suspension by two washes with Dulbecco’s Phosphate-Buffered Saline (DPBS) without calcium or magnesium, followed by treatment with 0.05% trypsin-EDTA. For fluorescent labeling of INS-1E cells, 1 mL of pre-warmed complete culture medium was added, and the cells were centrifuged for 5 minutes at 100 × g and 22°C. The medium was discarded, and the cells were resuspended in 1 mL of fresh culture medium. Hoechst 33342 fluorescent dye, diluted in DPBS to a final concentration of 20 μM, was added to the cell suspension at a 1:10 (v/v) ratio. Cells were incubated with Hoechst 33342 for 15 minutes at 37°C and 5% CO₂. The cell suspension was gently centrifuged for 5 minutes at 100 × g and 22°C, washed twice with DPBS, and gently resuspended in a small volume (20–30 μL) of fresh culture medium.

INS-1E cell suspensions (with or without fluorescent staining) from a single 10 cm culture dish were centrifuged for 1 minute at 100 × g and 22°C. The supernatant was removed, and 4 μL of the pellet was resuspended in 400 μL of Dulbecco’s Modified Eagle Medium (DMEM).

### Cell suspension freezing

A 1.5 μL aliquot containing approximately 80,000 cells was applied to 3 mm high-pressure freezing (HPF) planchettes with a 100 μm cavity (Leica Microsystems). A flat-faced planchette coated with 1-hexadecane was placed on top, excess medium was removed using filter paper, and the sample was vitrified using a Leica EM ICE HPF instrument (Leica Microsystems). After vitrification, the planchettes were separated in liquid nitrogen (LN₂), and the planchette with the 100 μm cavity was stored in LN₂ until further use. The planchette was transferred to FIB-SEM microscope.

### Cos-7 culture cultivation and freezing

Cos-7 cells (CRL-1651, ATCC) were cultured in plastic culture dishes (Sigma Aldrich) to 90% confluency in DMEM supplemented with 10% fetal bovine serum (FBS; Sigma Aldrich). Before cell seeding, plasma-cleaned TEM grids (Au, 200 mesh, R2/1, Quantifoil) were placed into cell culture dishes (Nunc Lab-Tek, Thermo Fisher Scientific) and UV-sterilized for 20 minutes. Following sterilization, the grids were incubated in the same culture medium as described above at 37°C for 2 hours to promote cell adhesion. A cell suspension with a density of 1.25 × 10⁵ cells/mL was prepared, and 100 μL of this Cos-7 cell suspension was applied to each grid. The cells were incubated at 37°C in a humidified atmosphere containing 5% CO₂ for 16 hours. Before plunge freezing, the grids were washed twice with phosphate-buffered saline (PBS; Sigma Aldrich, D1408) to remove non-adherent cells and cellular debris.

### Staphylococcus aureus biofilm

For biofilm experiments, we used *Staphylococcus aureus* ISP479C rsbU⁺ strains cultivated in a microfluidic culture system, as previously described^1^. The biofilm grows inside a fluorinated ethylene propylene (FEP) tube with an inner diameter of 300 µm. The FEP tube was coated with gold (40 nm) on the outer side. The mature biofilm was infected by phi812 (GeneBank KJ206563) 10^9^ PFU/ml, and then the FEP tube with infected biofilm was cut into 3 mm pieces which were frozen in 6 mm carriers using high pressure freezer Leica EM ICE (Leica Microsystems) with polyvinylpyrrolidone (50%) as cryoprotectant. The frozen FEP tube was cut lengthwise with an ultra-diamond knife 45° in ultra-microtome Leica UC7 (Leica Microsystems) under cryo-condition (-160°C) and then subjected to scanning electron microscopy.

### Tau protein fibrils

HEK293T Tau RD P301S FRET biosensor cells (ATCC-CRL-3275 provided by Dr. Monika Žilková, Institute of Neuroimmunology, Slovak Academy of Sciences) express Tau RD (244-368) tagged with either CFP or YFP. Upon seeding with proteopathic Tau seeds, endogenous Tau RD-CFP and Tau RD-YFP form filamentous Tau protein aggregates, generating FRET signal^2^. In our case, the proteopathic Tau seeds were sonicated The recombinant dGAE (Tau fragment 297-391) fibrils (<200 nm length) were sonicated and subsequently used for seeding the proteopathic Tau in this work.

dGAE (Tau 279-391) in PET17b plasmid in BL21(DE3) cells was expressed in LB medium and purified by two rounds of cation exchange chromatography at HiTrap SP HP (Cytiva) with buffer A (50 mM PIPES, 1 mM EDTA, 2 mM DTT, 0.1 mM PMSF, pH 6.9) and gradient of buffer B (A plus 1 M NaCl) followed by size exclusion chromatography on HiLoad Superdex 75 (Cytiva) with PBS pH 7.4. dGAE fibrils were produced by aggregation of dGAE (4 mg/ml) dialyzed into 10 mM sodium phosphate buffer, pH 7.2, with 130 mM NaCl and 10 mM DTT agitated in 96-well polystyrene PerkinElmer Isoplates for 16-48 hours at 200 rpm (double-orbital shaking, ClarioSTAR Plus) at 37°C. dGAE fibrils were sonicated on ice using a 1/16’’ microtip probe (Sonicator Q700, Qsonica) for 1.5 minutes at 20% amplitude, with 5-second on/off pulses. The quality and size of dGAE fibrils and seeds were checked by negative stain EM. After sonication, Tau seeds were < 200 nm, a size suitable for seeding.

HEK293T Tau RD P301S FRET biosensor cells were grown in DMEM (+ 1x GlutaMAX + 4,5 g/l D-glucose + Pyruvate, Gibco), supplemented with 10% FBS (Biosera) and 100 U/ml penicillin + 100 μg/ml streptomycin (PAA Laboratories) in a 6-well plate and volume 1.5 ml. After 3 days of on-plate cultivation, cells were transduced with Tau seeds (final concentration 4.5 µg/ml) using Lipofectamine^TM^ 2000 reagent (Invitrogen – Thermo Fisher Scientific). Tau-lipid transduction complexes were prepared by mixing 1 μl Lipofectamine^TM^ 2000 + 25 μl Opti-MEM (Gibco) and 664 pmol of Tau seeds diluted in 25 μl Opti-MEM; incubated at room temperature for 15 min. After 3 days, cells were detached from the plate using Acutase^TM^ Solution (EMD Millipore Corp.), harvested cells were centrifuged, resuspended, and seeded onto fibronectin-coated (80 ug/ml, Human plasma-derived Fibronectin protein, 1918-FN, R&D Systems) grids (gold-sputtered Quantifoil R 2/4 Au 200 mesh) placed in Lab-Tek 16-well chamber slide (Thermo Fisher Scientific) at a density of 30000 cells per grid in 80 μl/well. Cells on grids were incubated for 1 day. Before vitrification, the cells’ nuclei were stained with 1 μg/ml Hoechst 33342 Solution (Thermo Fisher Scientific) for 10 min. The remaining non-internalised dye and Tau protein were washed from the cell surface by 2 min of incubation in 0.06% Trypsin in PBS. Finally, cells were incubated for 10 min in cryoprotectant (DMEM + 10% glycerol). Cells on grids were vitrified in liquid ethane by Vitrobot IV (Thermo Fisher Scientific). Grids were clipped into autogrids and stored in liquid nitrogen.

Semi-automated FIB micromachining of cellular lamellae was carried out using Arctis Cryo Plasma-FIB (Thermo Fisher Scientific). Grid was sputter-coated with platinum by micro-sputter followed by 350 nm thick organo-metalic coating using the GIS, followed by another layer of platinum. Regions of interest were selected using FRET signal of Tau aggregates in the biosensor cells. CLEM data were collected using iFLM, Tau FRET signal was detected at GFP and Hoechst-stained nuclei at DAPI setting, respectively. Automated milling was done at a 15° angle relative to the grid using Xe plasma at 30 kV. Three rough milling steps were performed at decreasing currents of 1.0 nA, 0.3 nA and 30 pA with patterns placed progressively closer to the selected lamella, followed by polishing at 30 pA with the final lamellae thickness set to 200 nm.

Tomographic tilt series were collected using a Titan Krios (Thermo Fisher Scientific) operated at 300 kV. Imaging was performed with a Gatan Bioquantum K3 direct electron detector in zero-loss mode, using a 10 eV energy filter slit. Tilt series were collected at pixel size 3.4 Å with - 15 μm defocus. Data were collected using SerialEM^3^ with a dose-symmetric tilt scheme^4^, an angular range of ±50° with 2.5° increments. The total electron dose was approximately 120 e^−^/Å^2^. Individual tilt images were recorded as seven-frame movies in counting mode; frame alignment was carried out using MotionCor2^5^. Tomograms were reconstructed using AreTomo^6^ with 4x binning.

### Muscle tissue

The hamstring muscle of a 12-month-old mouse (hetreozygous zQ175; B6.12951-Htt<tm1Mfc<190JChdi) was dissected immediately post-mortem and placed in Ringer’s solution (115 mM NaCl, 2.5 mM KCl, 1.8 mM CaCl_2_, 2.5 mM MgCl_2_ and 3 mM sodium phosphate buffer pH 7). Fascicles were mechanically separated. Samples were taken with a sample punch provided by Leica Microsystems (Wetzlar, Germany) and were placed on a flat specimen carrier (0.5-mm thick, 1.5 mm in diameter, 200-μm deep, Leica Microsystems, #16706898). A drop of a cryoprotectant (10% Ficoll) was placed on top and immediately blotted prior to high-pressure-freezing (HPF) in a Leica EMPACT2 device. Samples were maintained in liquid nitrogen until further processing for CVEM. All experiments complied with Spanish and European legislation.

### Cryo-FIB/SEM microscopy

All experiments (except those involving the gallium FIB) were conducted on Helios Hydra 5 CX (Thermo Scientific) plasma-FIB/SEM microscope. The instrument was equipped a four-gas plasma-FIB column (Xe, Ar, O, N), a cryo-stage, an integrated fluorescence light microscope (iFLM), micro-sputter with a platinum target, and two gas injection systems (GIS, one operating at 45 °C for room-temperature applications, and one operating at 28 °C for applications at cryogenic temperatures) for organometallic platinum layer deposition. Sample milling with all plasma ion sources was carried out at 30 kV. Samples were positioned into the HPF/autogrid sample shuttle with a 35° pre-tilt before loading to the microscope. The sample surface contaminations were purged away by FIB imaging (Xe, 0.3 nA) under shallow angle (5-10°). The sample surface was sputter coated with platinum utilizing an integrated microsputter with xenon ions (16kV, 99nA, 6 minutes) striking at a platinum target. Unless stated otherwise, cryo-SEM imaging was conducted using the immersion mode with a 1 kV electron beam, an imaging current of 25 pA, a pixel size of 2.5 nm, a dwell time of 25 ns, and 64x line integration. A through-lens detector (TLD, suction tube: +70 V, mirror: -15 V) was utilized for the secondary electron detection. Experiments requiring gallium FIB milling were conducted on a Helios 5 UX (Thermo Scientific) instrument equipped with a cryo-stage and a gallium ‘Phoenix’ ion column.

### FIB milling procedure

After sputter coating the sample surface with platinum conductivity layer, cell distribution was investigated using iFLM and suitable regions were selected for initial sample surface cleaning. The surface grazing was performed with Xe^+^ (0.3 nA) at 3° milling angle corresponding to the stage tilt 0° and stage rotation 180° on the system used in this study. Then, the sample shuttle was unloaded from the microscope, the grid or the HPF carrier were manually rotated by 90°, and the shuttle was loaded back to the microscope. Next, a fiducial marker was milled into the sample and the data acquisition position was precisely located using correlation of FM and FIB images. The image correlation was performed in MAPS (Thermo Scientific). Subsequently, the initial trench was milled at a 74° milling angle (stage tilt to 1° and stage rotation to 0°). The initial trench was milled with Xe^+^ ions with dimensions approximately 80 x 50 x 1 µm using a milling template file for silicon. Declared depth 1 µm in silicon template excavated approximately 50 µm of ice. The trench served as a milling face. In the next step, a micrometer-thick protective organometallic platinum layer was deposited from the GIS system. Then, the sample shuttle was oriented to 16° milling position (stage tilt to 13° and stage rotation to 180°) and the residual GIS was ablated from the surface of the imaging face (Xe^+^, 4 – 15 nA). The imaging face polishing was carried out under the same geometrical conditions with O^+^ ions and 0.61 nA. Finally, the stage for tilted to 51° (stage rotation 180°) for perpendicular imaging. The last two steps were iteratively repeated to acquire CVEM data.

### Micrograph resolution analysis

Single image Fourier ring correlation (siFRC) analysis was adapted as described in Rieger et al.^23^ and used for resolution estimation across the image. The image was first divided into 256×256 pixel patches, and the resolution was calculated for each patch as the value at which siFRC first dropped below 0.143. The final resolution was determined as the minimal value among all patches.

### Contrast estimation

The image was first low-pass filtered using a Gaussian blur (kernel size = 10, σ = 2). A minimum of five mitochondria were segmented, along with the same area of the cytoplasm filling the intercellular space. The Microscopy Image Browser v. 2.9010 (MIB)^42^ was used for semi-manual segmentation. Contrast was calculated using the Weber contrast metric^43^ implemented in Python script.

### Level of charging artifacts measurement

Thresholding of charging artifacts was used to determine the fraction of an organelle’s image affected by charging. The micrographs were manually inspected, and thresholds for reliable segmentation of charging artifacts were identified. If the estimated threshold produced only continuous regions corresponding to charging artifacts, the threshold value was increased. Once the mask began to include isolated regions outside the centers of charging artifacts, the procedure was stopped, and the previous threshold value was used to generate the final mask, which defined the fraction of the image affected by charging. This method was also used to determine the level of beam-induced damage.

### Analysis of milling speed and artifacts

Measurement of milling artifacts (“curtaining”) was carried out on high-pressure–frozen DMEM medium. FIB milling was performed at a 90° milling angle followed by SEM imaging at 38°. Three current levels were selected for each plasma ion source. At least ten images were automatically collected for the low (up to 1 nA) and mid-current (1 – 10 nA) settings, whereas six images were manually acquired for the high-current (more than 10 nA) settings. Imaging parameters were set to 1 kV acceleration voltage, 12.5 pA imaging current, 5 nm pixel size, and 25 ns dwell time with 128× line integration. All images obtained after plasma milling were cropped to 2700 × 2700 pixels.

Subsequently, the images were analyzed using an in-house Python-based algorithm. The algorithm highlights a signal containing vertical frequencies located within a 5° vertical wedge of the 2D Fourier spectrum. The resulting image contains only signal corresponding to vertical frequencies. The curtaining score is defined as the standard deviation of pixel values in this resulting image, with higher scores indicating stronger curtaining. The calculation is based on the method of M. Dumoux et al.^19^

Ice-milling rates were measured using vitrified DMEM. Organometallic platinum milling rates were determined using an approximately 5-µm-thick platinum layer deposited on vitrified DMEM. The proposed workflow was used to maintain perpendicular milling geometry. Trenches were milled using the cleaning-cross-section pattern with dimensions of either 7 × 3 µm or 3 × 3 µm. Milling was performed at a 16° angle. Low-current measurements used currents in the range of 20 pA to 2.4 nA, while high-current measurements used 15–200 nA. Three milling currents were selected within each of these ranges, and milling time was adjusted to maintain a constant final dose. The depth of the ablated trench was then measured from SEM images collected perpendicular to the milling direction. The final milling rate was calculated from the total applied charge and the volume of material removed.

### Monte-Carlo simulation

The Monte Carlo simulation method was employed to investigate the interaction of electrons with low-density ice (LDA). LDA with a density of 0.94 g/cm^3^ was used for the simulation. The simulation was conducted using CASINO v2.51 software using Mott interpolation. Two different electron beam energies, namely 1 kV and 5 kV, were simulated under zero tilt conditions (perpendicular imaging).

### Autofocus algorithm

The autofocus algorithm sweeps the optimized variable (such as working distance - WD) during a single scan in a serpentine pattern. For each scanned line, a focusing criterion is calculated, corresponding to a specific WD. The criterion is based on power spectral density in middle frequencies. Given that the sweeping process is repeated multiple times within a single scan, the focusing criterion is averaged across lines that correspond to the same WD value. The WD that corresponds to the best focusing criterion is identified, and the corresponding WD value is used. The same routine has been implemented to reduce image astigmatism and centering the beam to optical axis. Concurrently, a focus index criterion^25^ was implemented as an alternative focusing criterion. These functions were combined with in-line image auto-optimization described in Xu et al.^25^ and a template matching algorithm for drift correction.

### Elastic alignment of cryo-data

Rigid alignment of consecutive slices was performed using SIFT^26^ as implemented in Fiji^44^ on the raw data with a voxel size of 2.5 nm × 2.5 nm × 10 nm. The following settings were used for SIFT-based alignment: initial Gaussian blur: 0.5 px; steps per scale octave: 3; minimum image size: 64 px; maximum image size: 1024 px; feature descriptor size: 8; feature descriptor orientation bins: 16; closest/next-closest ratio: 0.92; maximal alignment error: 25 px; inlier ratio: 0.05; expected transformation: translation. The maximal alignment error was later increased to 70 px to accommodate larger expected shifts between keypoints in the frames. The voltage-contrast nature of CVEM imaging introduces additional deformations. Images obtained using the presented method were observed to be susceptible to line-to-line signal shifts. The intensity values within each scanned line are correctly positioned relative to one another; however, a displacement occurs between adjacent lines. These displacements are correlated within a single slice, whereas the displacement of corresponding lines between consecutive slices is uncorrelated. The deformation can therefore be modeled as a two-dimensional displacement field, with one shift value assigned to each line within the volume. The displacement field is determined by calculating a running average with a window size of 10 lines within each slice. This produces a smooth displacement field appropriate for the low signal-to-noise data and is consistent with the expected deformation behavior. Subsequently, a pairwise cross-correlation is computed between corresponding lines in consecutive slides. To improve computational efficiency, only shifts smaller than a specified threshold (60 px) are considered. In the next step, the true line positions are recovered by calculating the mean displacement across five frames, leveraging the independence of line shifts between slices. Finally, the calculated displacements are applied to the raw images.

### Volume segmentation

Data was down-sampled to the isotropic voxel size of 20 nm before automated segmentation. Raw data was normalized using quantile normalization between quantiles 0 and 0.99. Manual ground truth was done using Napari^45^ for the following classes: foreground, cell boundary, lipid droplets, mitochondria and nucleus boundary. 3D U-Net was trained using the Monai framework (https://monai.io/).The network consisted of 3 resolution levels with 32, 64 and 128 feature maps each. Each level had two convolutional blocks with BatchNorm normalization. The networks were trained on NVIDIA GeForce RTX 3090 with patch size of 128×128×128 and batch size 2 for 15 epochs of 1200 samples using Adam optimizer with learning rate 0.0001 and weight decay 0.0005. Random augmentations including flip, gaussian blur with sigma from 0.6 to 2, intensity scaling by a factor between -0.2 and 0.2 and intensity shifting with an offset between - 0.1 and 0.1 were applied with 0.5 probability. Low contrast of the data, small amount of ground truth and sparsity of features in the volume made it necessary to adjust the common U-Net training procedure. The sampling of training examples was biased towards the positive class (70% of samples in training contained the segmented class). After the foreground of the cell was segmented, for all other classes the masked Dice loss was used to ignore the gradients from the background pixels. After 15 epochs the accumulated statistics in the normalization layers were fixed and the networks were trained for another 15 epochs with learning rate 0.00001 to compensate for the instability of the batch statistics resulting from the small batch size.

### Sub-volume averaging of Cos-7 nuclear pore complexes

The overall number of 239 NPC volumes was manually selected from a cryo-volume data of single Cos-7 cell. The individual NPCs were extracted with 128×128×39 voxels (voxel size 3.05×3.05×10 Å), rescaled to uniform 3.05 Å voxel size using bilinear interpolation, following by contrast inversion, normalization of intensities, and finally the volumes were cropped to 96×96×96 voxels for the subsequent processing. The sub-volume averaging was carried out according to gold-standard Fourier shell correlation (FSC) protocol. The two initial models were created by independent phase randomization of the cryo-EM map of *S. pombae* NPC^46^ (EMD-11373) above 200 Å resolution. Subsequently, the reference volumes were bandpass filtered to 1000-200 Å. Twelve iterations of global volume alignment with angular step of 15° and 7.5° and maximum shift of +/-5 voxels was carried out independently for each half-set. The normalized cross-correlation coefficient (ncc) was used to evaluate NPC alignment to the reference. The new reference calculated from the particles after each iteration was bandpass filtered to 1000-200 Å before entering next iteration. The NPC volume with unstable alignment parameters in last four iterations or ncc<0.1 were excluded from further analysis leaving 113 particles for the next refinement steps. The NPC volumes were then subjected to five iterations of global refinement (angular step 7.5°) followed by five iterations of local refinement. At this stage, C8 symmetry was imposed during reconstruction and the lowpass filtering of the initial was gradually changed from 200 Å to 70 Å during this iteration cycle. The Cos-7 NPC density was refined to 94 Å (FSC_0.143_) using this approach. The sub-volume picking was carried out in Napari (cit. Napari) and all the volume manipulation, alignment, and reconstruction steps were carried out using an in-house written code implemented in Python.

## Data availability

The raw CVEM data for cos-7 cell, INS-1E cell, rat muscle tissue are available for download from https://aperture.ceitec.muni.cz/cryoem/experiments/0198e6ea-0dfd-7b6c-bc6e-1fc6175df5d5

## Code availability

The graphical user interface program for acquisition of CVEM data is available at https://github.com/cemcof/FIBSEM_Maestro. The software tool for the correction of line-by-line deformation is available at https://github.com/kreshuklab/cryofib_crosscorr. The algorithm for sub-volume averaging is available at https://github.com/cemcof/cvem_sva.

## Acknowledgements

The work was supported by the Czech Science Foundation (22-02203S to JN, 22-15175I to HN). We acknowledge Cryo-electron microscopy and tomography core facility CEITEC MU of CIISB, Instruct-CZ Centre, supported by MEYS CR (LM2023042) and European Regional Development Fund-Project „Innovation of Czech Infrastructure for Integrative Structural Biology“ (No. CZ.02.01.01/00/23_015/54). This work was supported by the project National Institute for Research of Metabolic and Cardiovascular Diseases (Program EXCELES, ID Project No. LX22NPO5104), funded by the European Union-Next Generation EU) and by the National Institute of Virology and Bacteriology (Program EXCELES, ID Project No. LX22NPO5103) funded by the European Union - Next Generation EU. AK and PK acknowledges the support from EU through HEU program (IMAGINE project, grant agreement 101094250-IMAGINE). EB was supported by the Joachim Herz Foundation through an Add-on Fellowship for Interdisciplinary Life Science. The support from the project New Technologies for Translational Research in Pharmaceutical Sciences /NETPHARM, project ID OP JAC CZ.02.01.01/00/22_008/0004607 co-funded by the European Union is acknowledged. The work was supported by grant PID2023-153013OB-I00 funded by MCIU/AEI/10.13039/501100011033/FEDER, UE) awarded to MRF-F. JN acknowledges funding from the NOMIS Foundation.

## Author contributions

PK: conceptualization, investigation, methodology, data analysis, manuscript preparation and review; JM: investigation, methodology; ZT: investigation, methodology; EB: methodology; PP: supervision; AK: supervision; JN: conceptualization, methodology, supervision, manuscript preparation, writing, review.

## Supporting information

**Figure S1:**
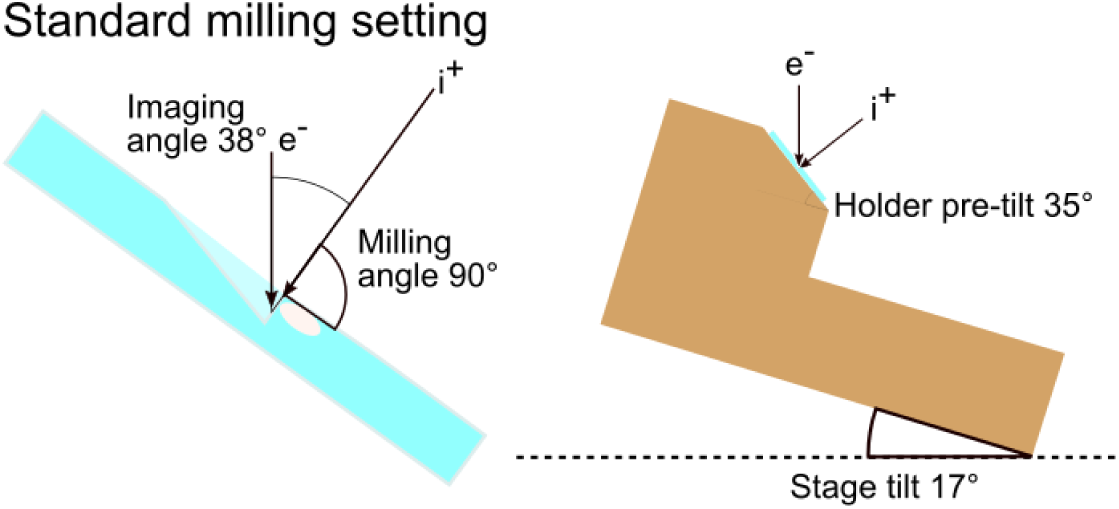
Geometry of the “through-the-trench” FIB/SEM experiment, in which ion beam milling is performed in perpendicular direction to the sample surface and SEM imaging is carried out under an angle 38° angle(valid for FIB/SEM instruments with 52° angle between electron and ion beam column).

**Figure S2:**
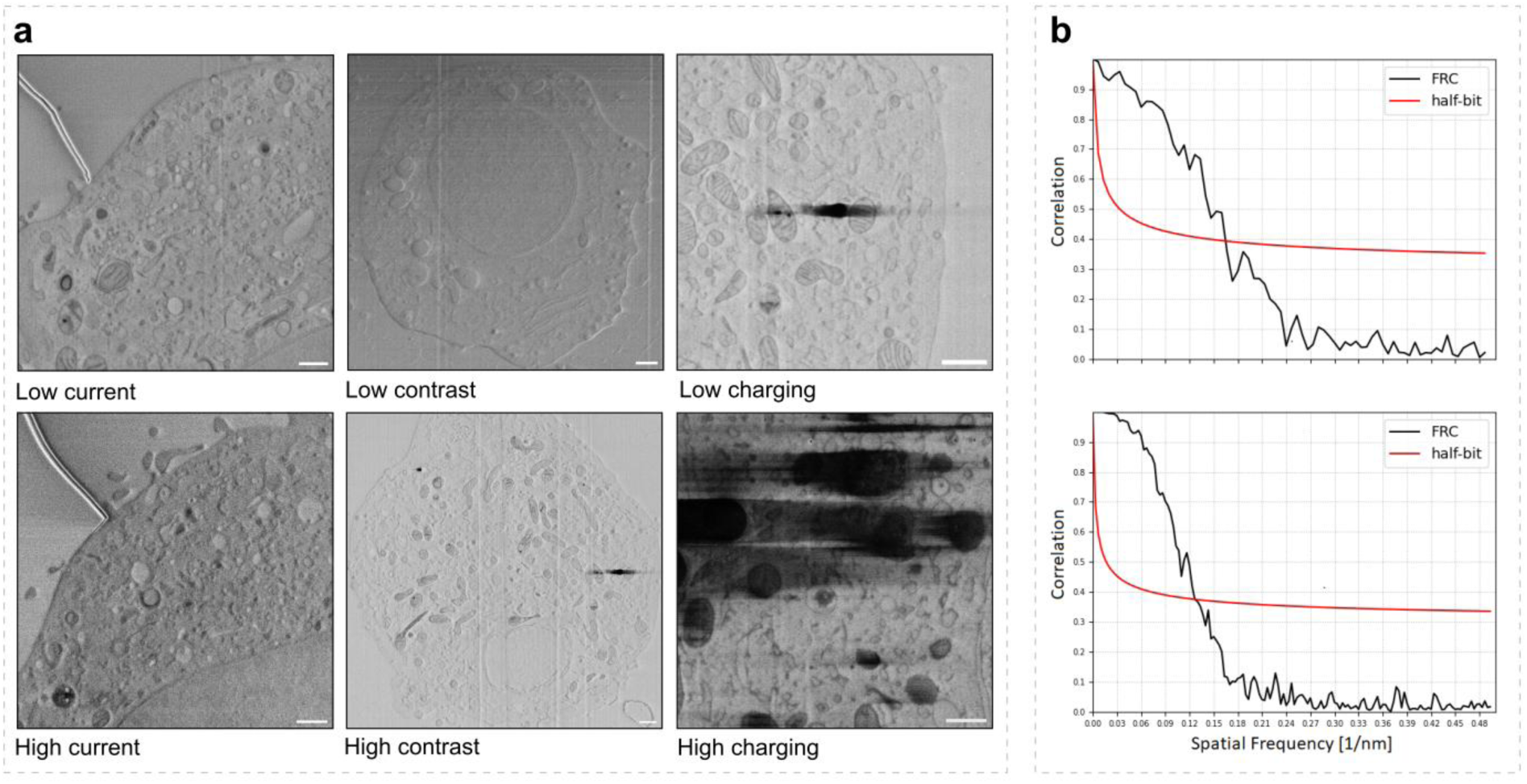
Comparison of images acquired under with low and high imaging current; conditions resulting in low and high contrast; and conditions leading to low and high content of charging artifacts (A). The scale bars correspond to 200 nm. A siFRC curve for and image acquired under through-the-trench geometry (B top) and perpendicular imaging (B bottom), respectively.

**Figure S3:**
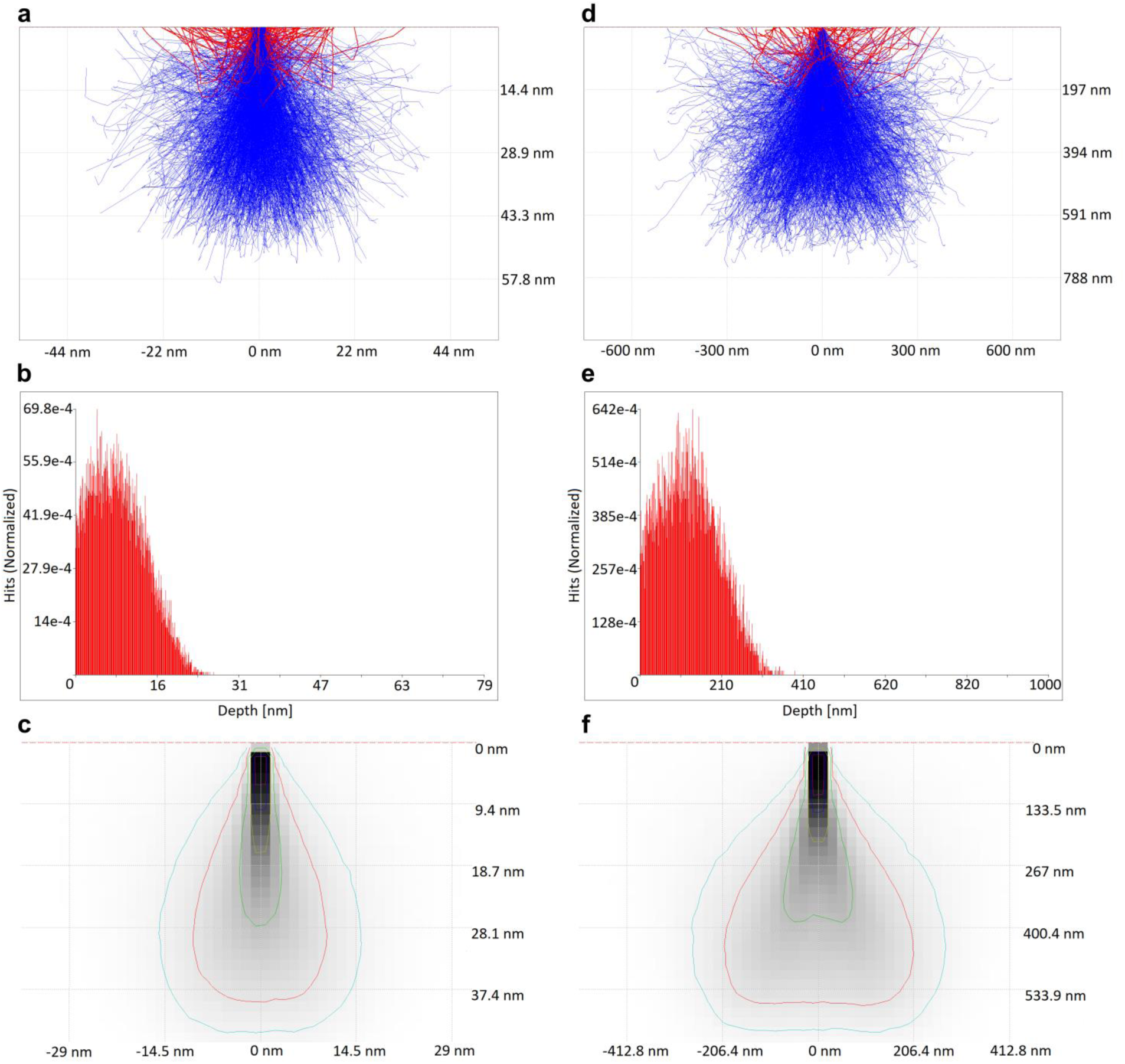
Results of Monte Carlo simulations for 1 keV primary electron beam energies (A, B, C) and 5 keV electron beam energies (D, E, F), respectively. Electron scattering trajectories for secondary electrons (blue) and back-scattered electrons (red; A,D). Maximum penetration depth of electron beam trajectories, which eventually escape from the sample (B, E). Cross-section plot of absorbed energy (C, F). Light blue line corresponds to 5% of energy, red to 10%, green to 25%, yellow to 50%, dark blue to 75%, and purple to 90% of energy.

**Figure S4:**
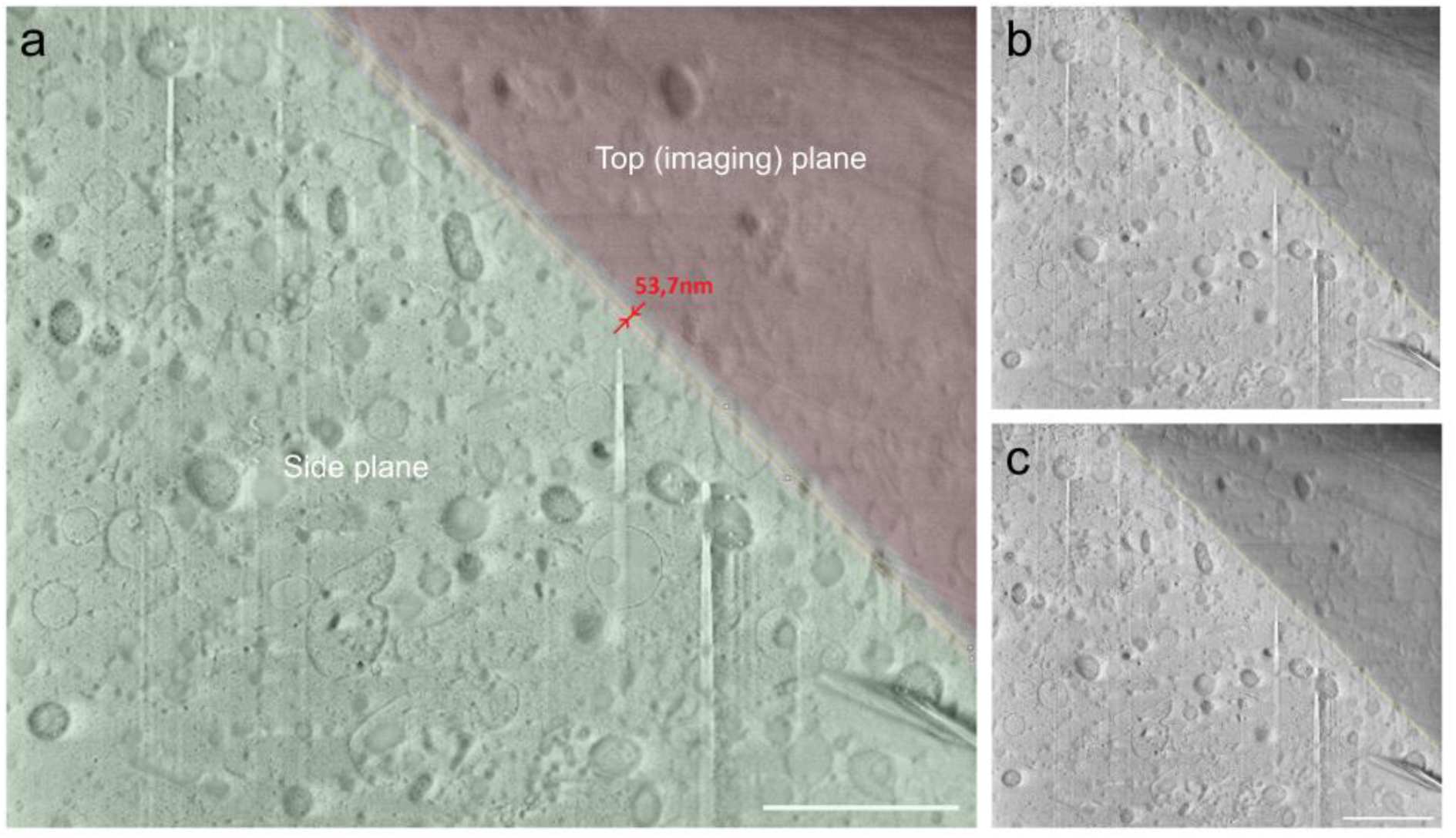
Verification of the sample sectioning precision. An orthogonal view on the imaging face from the side trench was acquired before and after ablation of five 10 nm sections (A). An image of the side face at the beginning of the experiment. Image of the side face after ablation of five 10 nm sections. The scale bars correspond to 1 μm.

**Figure S5:**
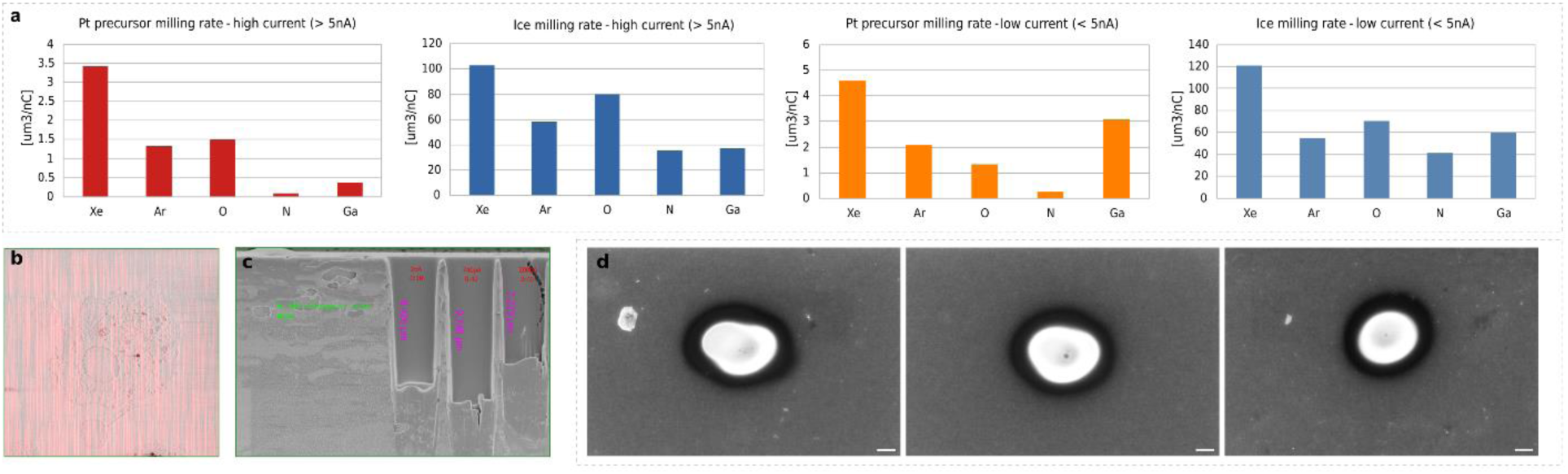
Milling rates of platinum precursor and vitrified ice by different ion sources. The plots are further divided into milling by low (<5 nA) and high (>5 nA; C, D) currents, respectively. An example image highlighting segmentation of curtains by FFT filtering for the curtaining score calculation (B). Milling of 7 x 3 μm trench into platinum precursor with 30 kV Ar^+^ beam with 2 nA (t=18 s; *left*), 740 pA (t=49 s; *middle*), 200 pA (t=180 s; *right*) results in trenches with a depth of 8.4 μm, 9.7 μm, and 7.2 μm, respectively (C). Spot burns using 240 nA O+ plasma (30 kV) into silicon wafer after 5 s milling. The images show snapshots from the oxygen probe alignment according to protocol presented here. The profile of non-compensated beam (*left*), partially aligned beam (*middle*), and the shape of fully compensated probe (*right*) are shown. Scale bars correspond to 200 nm.

**Figure S6:**
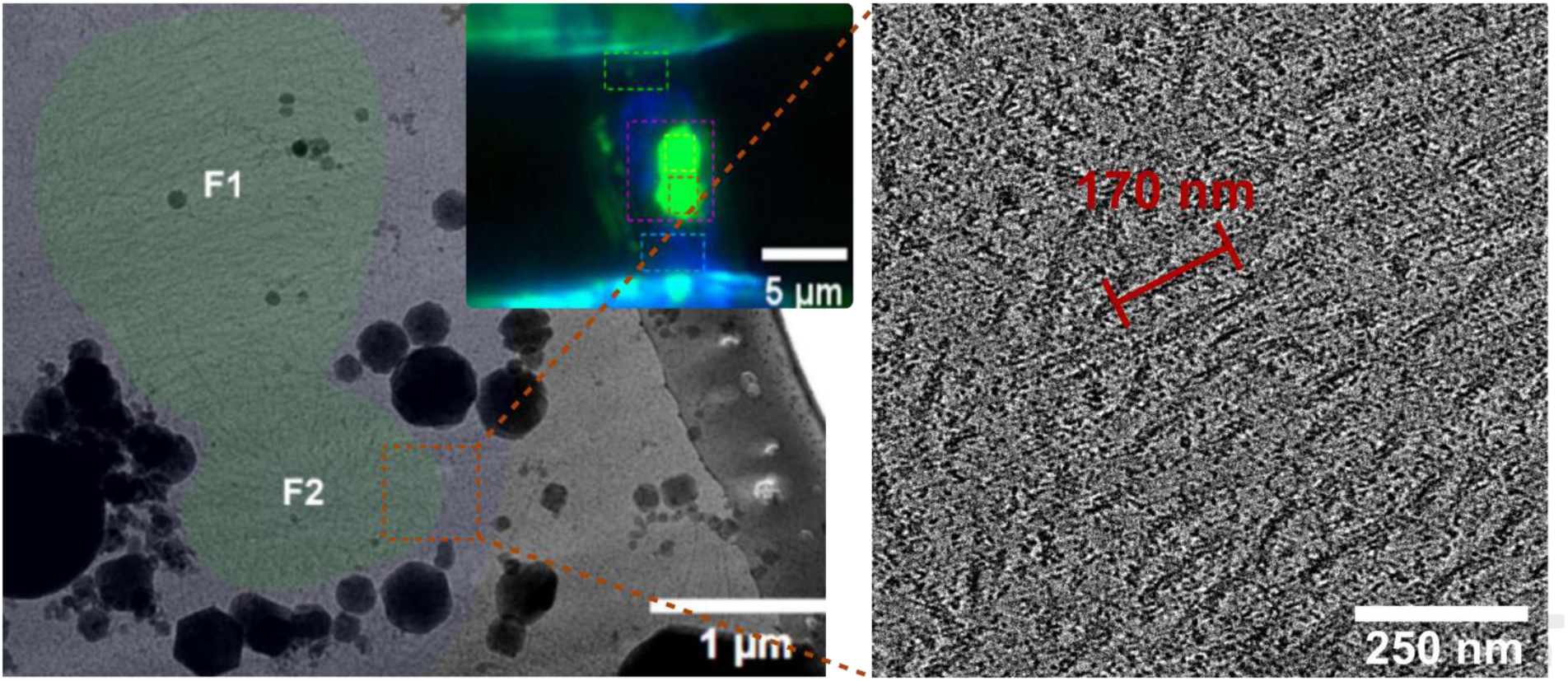
Cryo-TEM (cryo-FM in inset) image of HEK293T lamella showing filamentous tau (green) and a slice from cryo-ET tomogram highlighting measured filament pitch.

